# Assessment of two DNA extraction kits for profiling poultry respiratory microbiota from multiple sample types

**DOI:** 10.1101/2020.10.21.348508

**Authors:** Michael E.C. Abundo, John M. Ngunjiri, Kara J.M. Taylor, Hana Ji, Amir Ghorbani, KC Mahesh, Bonnie P. Weber, Timothy J. Johnson, Chang-Won Lee

## Abstract

Characterization of poultry microbiota is becoming increasingly important due to the growing need for microbiome-based interventions to improve poultry health and production performance. However, the lack of standardized protocols for sampling, sample processing, DNA extraction, sequencing, and bioinformatic analysis can hinder data comparison between studies. Here, we investigated how the DNA extraction process affects microbial community compositions and diversity metrics in different chicken respiratory sample types including choanal and tracheal swabs, nasal cavity and tracheal washes, and lower respiratory lavage. We did a side-by-side comparison of the performances of Qiagen DNeasy blood and tissue (BT) and ZymoBIOMICS DNA Miniprep (ZB) kits. In general, samples extracted with the BT kit yielded higher concentrations of total DNA while those extracted with the ZB kit contained higher numbers of bacterial 16S rRNA gene copies per unit volume. Therefore, the samples were normalized to equal amounts of 16S rRNA gene copies prior to sequencing. For each sample type, all predominant taxa detected in samples extracted with one kit were present in replicate samples extracted with the other kit and did not show significant differences at the class level. Furthermore, between-kit differences in alpha and beta diversity metrics were statistically indistinguishable. Therefore, both kits perform similarly with regard to 16S rRNA gene-based poultry microbiome analysis.

## Introduction

The importance of gut and respiratory microbiota in poultry health and production performance has resulted in an increase of studies aimed towards developing microbiome-based intervention strategies [1–5]. Consequently, different protocols for sampling, sample processing, DNA extraction, sequencing, and bioinformatic analysis have emerged. The varying protocols utilized by different research groups can introduce confounding factors and complicate data comparison between studies. For example, protocol-associated-between-studies-variation in biodiversity profiles in human gut and environmental microbiota has been widely reported [6–14]. We and others have evaluated the sample collection methods for poultry respiratory and gut microbiota [2, 15–17]. However, protocol-associated-confounding-factors for DNA extraction, sequencing strategies, and bioinformatic analysis in poultry microbiota remain to be studied.

An investigation carried out by the Microbiome Quality Control Project, where identical sets of samples were processed in 15 laboratories, identified the DNA extraction process as a significant cause of variation between laboratories, only second to sample type and origin [7]. Efficient and bias-free bacterial DNA extraction for microbiota analysis requires effective disruption of the diverse cell wall structures and compositions present in different types of bacteria [18]. Kits for bacterial DNA extraction are designed to disrupt cell walls mostly through mechanical, chemical, and enzymatic methods [19]. Each of these lysis methods has its own advantages and disadvantages. While mechanical lysis can disrupt all types of bacteria and increase DNA yield, it can shear or fragment genomic DNA and compromise the outcomes of downstream applications, including sequencing [20–21]. Enzymatic and chemical lysis methods are less likely to damage DNA, but their inefficiency in disrupting certain types of bacteria may introduce bias in the biodiversity profiles [22]. Another important consideration for choosing extraction kits are the contaminating DNAs that are ubiquitously present in kits and reagents. Kit contaminants, especially nucleotide contaminants, critically impact the biodiversity results obtained from low biomass samples [23–24].

Five different DNA extraction kits have been used in poultry respiratory microbiota studies, including BiOstic FFPE Tissue DNA Isolation Kit, PowerSoil DNA Isolation Kit, QIAmp DNA Mini Kit, Maxwell RSC PureFood Pathogen Kit, and Qiagen DNeasy Blood and Tissue Kit [2, 15–17, 25–28]. These kits typically employ a combination of mechanical, chemical, and enzymatic disruption methods to increase DNA yields. In our previous poultry respiratory microbiota studies [2, 15–17], the Qiagen DNeasy Blood and Tissue kit (BT kit), which utilizes chemical and enzymatic lysis, was used. The BT kit uses a broad-spectrum serine protease (proteinase K) that can potentially lyse all types of bacteria, but an optional use of lysozyme can ensure complete lysis of Gram-positive bacteria. However, the BT kit is not certified for the extraction of low biomass samples, such as respiratory tract swabs, without the use of carrier nucleic acids, which can complicate downstream applications [29].

In this study, we compared the biodiversity profiles of chicken respiratory microbiota generated from DNA samples extracted using the BT kit and ZymoBIOMICS DNA Miniprep kit (ZB kit). The ZB kit uses a combination of mechanical and chemical disruption methods and is a certified low bioburden kit [30]. Pre-sequencing results showed great differences between the BT and ZB kits in DNA yield and 16S rRNA gene copies per unit volume. However, after sequencing of normalized samples, both kits produced similar microbiome data in terms of bacterial diversity and class-level taxonomic profiles.

## Materials and Methods

### Animals and ethics statement

Animal management and euthanasia were in correspondence with protocol #2015A00000056 approved by The Ohio State University Institutional Animal Care and Use Committee. This protocol conforms with the U.S Animal Welfare Act, Guide for Care and Use of Laboratory Animals and Public Health Service Policy on Humane Care and Use of Laboratory Animals. The Ohio State University is accredited by the Association for the Assessment and Accreditation of Laboratory Animal Care International. White leghorn chickens were obtained from our institution’s (Food Animal Health Research Program, Wooster, OH [17]. The chickens were provided with *ad libitum* access to feed and water and their welfare was monitored twice daily. All birds were clinically healthy and were all included in the study. Prior to the collection of invasive samples, the birds were humanely euthanized through carbon dioxide exposure as previously described [17].

### Experimental Design

A total of 72 four-week-old SPF chickens were housed in the same room and were randomly assigned to three groups for sampling (Fig 1). The sample size was decided based on our recent reports showing that four or more respiratory samples were sufficient for 16S based taxonomic and diversity analysis [17]. An assortment of invasive and non-invasive respiratory samples including, nasal cavity wash, upper and lower tracheal wash, lower respiratory lavage, as well as choanal and tracheal swabs were obtained from either live or euthanized birds as described in our recent study [17]. The samples were pooled as follows: 24 samples per pool for tracheal swabs from live birds, choanal swabs from live birds, tracheal swabs from euthanized birds, choanal swabs from euthanized birds, upper tracheal wash pellets, lower tracheal wash pellets; and 48 samples per pool for nasal cavity wash pellets and lower respiratory lavage pellets. The pooled samples were homogenized and divided into 8 replicates for tracheal and choanal swabs and tracheal wash pellets, and into 16 replicates for the nasal cavity wash pellets and lower respiratory lavage pellets. The sample pools were evenly divided to be extracted with either the BT or ZB kit. This study did not involve treatment groups and, therefore, no control animals were required.

**Fig 1.**
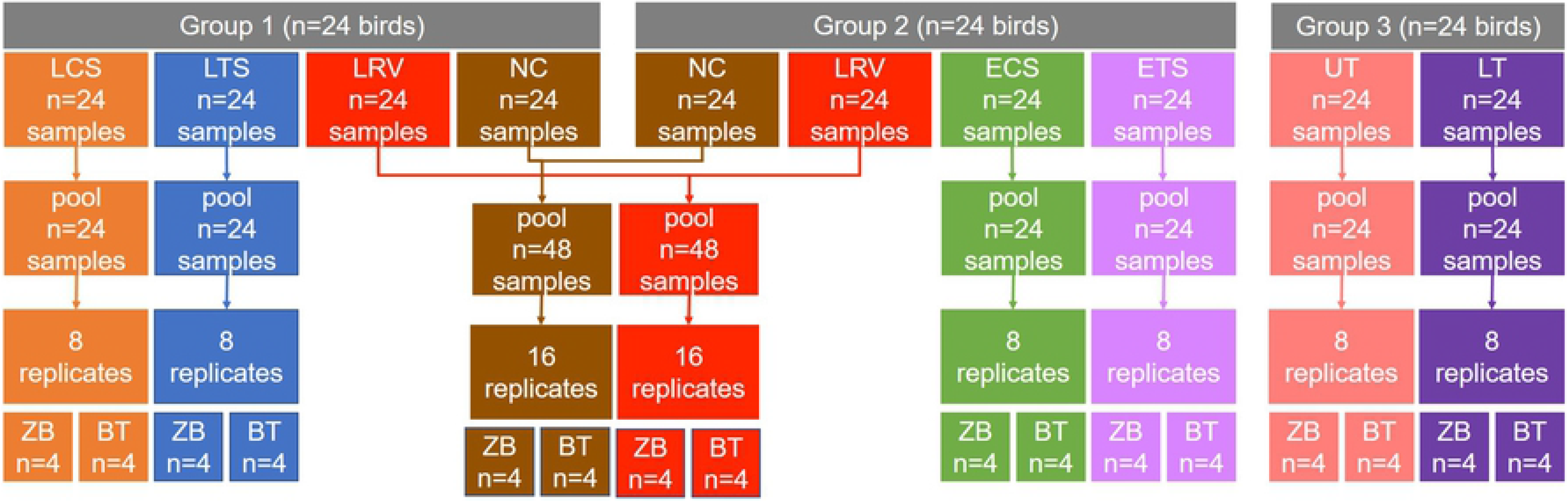
Experimental Design. A total of 72 four-week-old SPF chickens were divided into three groups. Samples were collected from individual birds and pooled as shown above. Each sample pool was aliquoted into several replicates which were evenly divided between the BT and ZB kits. LCS, live choanal swab; ECS, euthanized choanal swab; LTS, live tracheal swab; ETS, euthanized tracheal swab; NC, nasal cavity wash; UT, upper trachea wash; LT, lower trachea wash; LRV, lower respiratory lavage.

### Sample Processing and DNA extraction

Sample collection and processing were done as described in our recent report [17]. DNA extraction using the BT kit was previously described [17]. Extraction of DNA from respiratory samples using the ZB kit was performed per the manufacturer’s direction. In addition, the final elution volume for both the BT and ZB kits was standardized to 50 μL to allow a side-by-side comparison of DNA yields. Furthermore, both a mock community control (ZymoBIOMICS Microbial community standard) and negative extraction controls (1X PBS) were extracted alongside the samples using both the BT and ZB kits. Each negative extraction control was subjected to each processing step, all the way from sample collection to 16S rRNA sequencing.

### 16S rRNA gene sequencing, analysis, and sample filtering

Our optimized methods for pre-sequencing and sequence processing are detailed in our recent reports [16–17] and are stated here in brief. The 16S rRNA V4 hypervariable region was amplified using primer sets 515F (GTGYCAGCMGCCGCGGTAA) and 806R (GGACTACNVGGGTWTCTAAT). Samples were normalized to 1.67 × 10^5^ 16S rRNA gene copies/μL prior to barcoding and Illumina MiSeq sequencing (reagent kit v3; Illumina, San Diego, CA) at the University of Minnesota Genomics Center (Minneapolis, MN). Afterwards, sequences were demultiplexed, merged, and processed to remove primer sequences and sequences < 245 and > 260 base pairs. Additionally, sample- and taxonomic-based filtering methods were applied to the samples (see Results).

Further sequence processing, including denoising, amplicon sequence variant (ASVs) clustering, non-16S rRNA ASV removal, mid-rooted phylogenetic tree generation, and taxonomic classification through SILVA 16S rRNA gene database (release 132) were implemented in QIIME2 (qiime2-2019.1) [31]. Statistical differences between DNA quality and quantity, 16S rRNA gene copies in total DNA per sample, number of 16S rRNA gene sequences per sample, and alpha diversity were examined using a nonparametric Kruskal-Wallis test with Benjamini-Hochberg correction for false discovery [32]. Moreover, statistical differences in beta diversity were examined using a pairwise permutational multivariate analysis of variance (PERMANOVA) with a Benjamini-Hochberg correction for false discovery [33]. Differences were considered significant at p < 0.05.

### Data availability

Raw data files and metadata are publicly available in the NCBI BioProject database under accession number PRJNA668323 and can be accessed through https://www.ncbi.nlm.nih.gov/sra/PRJNA668323. Raw data files are also available through the Sequence Read Archive (SRA) under accession numbers SRX9277982 - SRX9278111.

## Results

### BT and ZB kits differ greatly in DNA yield and 16S rRNA gene copy numbers

For both kits, DNA was eluted in a volume of 50 μL to allow between-kits comparison of DNA concentrations and purity. DNA quality was spectrophotometrically measured and expressed as ratios of absorbances at 260nm and 280nm wavelengths (A260/A280) and 260nm and 230nm wavelengths (A260/A230). The mean A260/A280 values for most of the samples were around 1.8 and were not statistically different between the kits. However, the mean value for the lower respiratory lavage samples extracted with the ZB kit was higher than 2.0 which was significantly higher compared to the BT kit (Fig 2A). On the other hand, higher mean A260/A230 ratios were observed for most of the samples extracted using the BT kit, with significantly higher ratios in the live choanal swab, euthanized choanal swab, euthanized tracheal swab, nasal cavity wash, and lower respiratory lavage (Fig 2B). DNA yield was spectrophotometrically measured and expressed as mass per unit volume (ng/μL). In general, the BT kit tended to have more DNA yield than the ZB kit, with DNA quantities being significantly higher for most of the sample types except for the lower trachea wash and lower respiratory lavage (Fig 2C).

**Fig 2.**
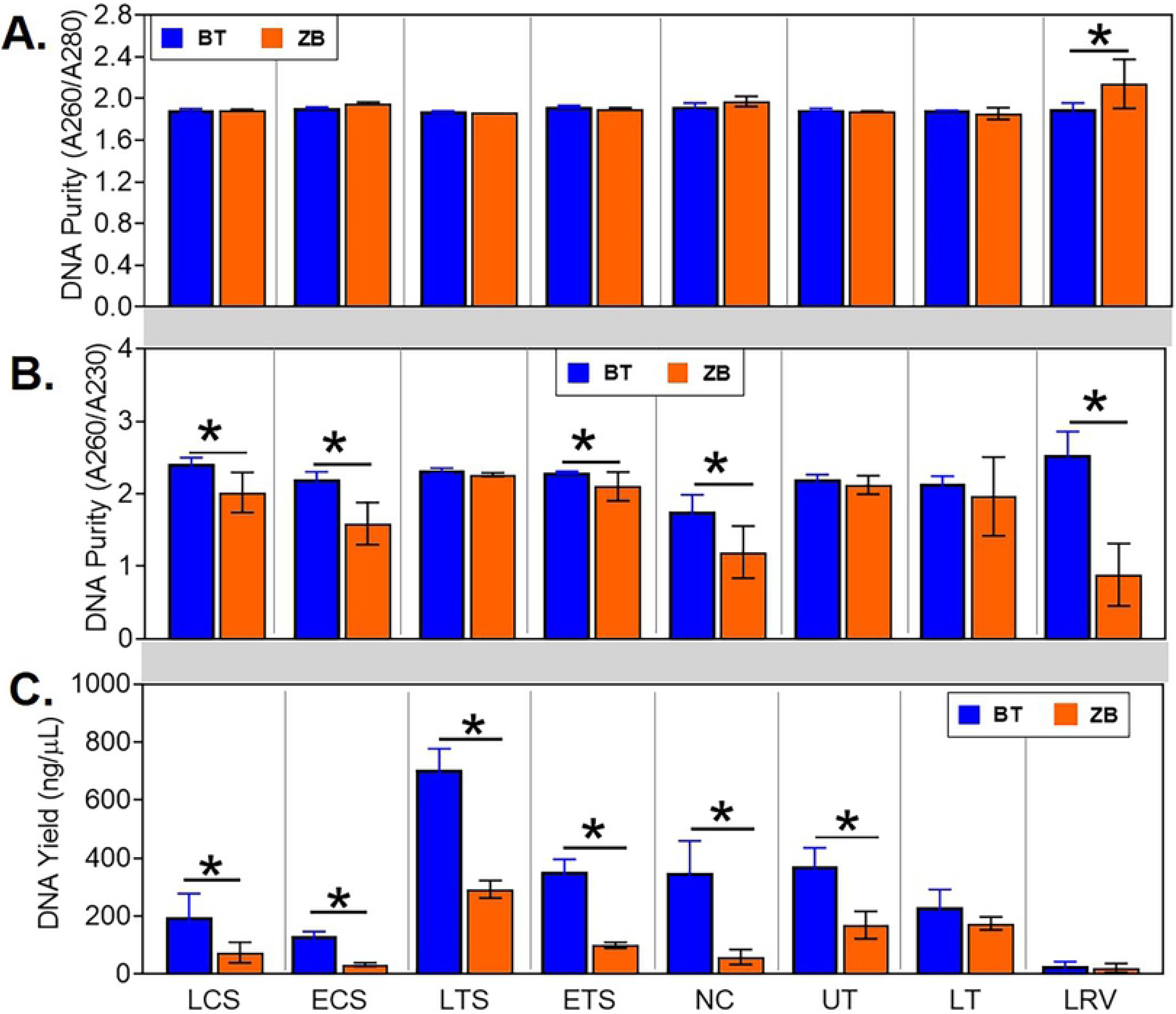
Comparison of quantity and quality of DNA from samples extracted using the ZB and BT kits. The elution volume was standardized to 50 μL for both kits. DNA quality was expressed as ratios of absorbances at (A) 260nm and 280nm wavelengths (A260/A280) and (B) 260nm and 230nm wavelengths (A260/A230). (C) DNA yield was expressed as mass per unit volume (ng/μL). Statistical differences, * p < 0.05 (pairwise Kruskal-Wallis tests, with a Benjamini-Hochberg correction for false discovery). LCS, live choanal swab; ECS, euthanized choanal swab; LTS, live tracheal swab; ETS, euthanized tracheal swab; NC, nasal cavity wash; UT, upper trachea wash; LT, lower trachea wash; LRV, lower respiratory lavage.

The DNA was subsequently subjected to quantitative PCR to determine the number of 16S rRNA gene copies. Negative extraction controls were included to facilitate filtering out of samples with little-to-no 16S rRNA gene copies. The highest amounts of 16S rRNA copies that were detected in the negative controls were set as elimination thresholds for the corresponding sample types. The elimination cut-offs for the BT kit were 1.6 × 10^3^, 1.4 × 10^4^, 2.0 × 10^3^, 3.1 × 10^3,^ and 1.1 × 10^4^ 16S rRNA copies per nanogram of total DNA for choanal swab, lower respiratory lavage, nasal cavity wash, tracheal swab, and lower and upper tracheal washes, respectively. The elimination cut-offs for the ZB kit were 20, 50, 20, 60, and 15 16S rRNA copies per nanogram of DNA for choanal swab, lower respiratory lavage, nasal cavity wash, tracheal swab, and tracheal wash, respectively. Using these criteria, 3 out of the 40 samples extracted using the BT kit were dropped from further analysis (1 live tracheal swab and 2 lower respiratory lavage samples). In contrast, all 40 samples extracted using the ZB kit were retained.

Among the retained samples, the nasal cavity wash, euthanized tracheal swab, and euthanized choanal swab samples extracted with the ZB kit had significantly higher numbers of 16S rRNA gene copies than those extracted using the BT kit (Fig 3A). Subsequently, all samples were normalized to 1.67 × 10^5^ 16S rRNA gene copies/μL. This was done to achieve 5 × 10^5^ 16S rRNA gene copies in 3μL of normalized sample used for pre-sequencing PCR, which approximately translates to a target sequencing coverage of at least 10X. The PCR products were subsequently used for sequencing library preparation. No significant differences in the 16S rRNA gene sequence counts from the normalized samples were observed between the kits, except for the lower respiratory lavage samples (Fig 3B).

**Fig 3.**
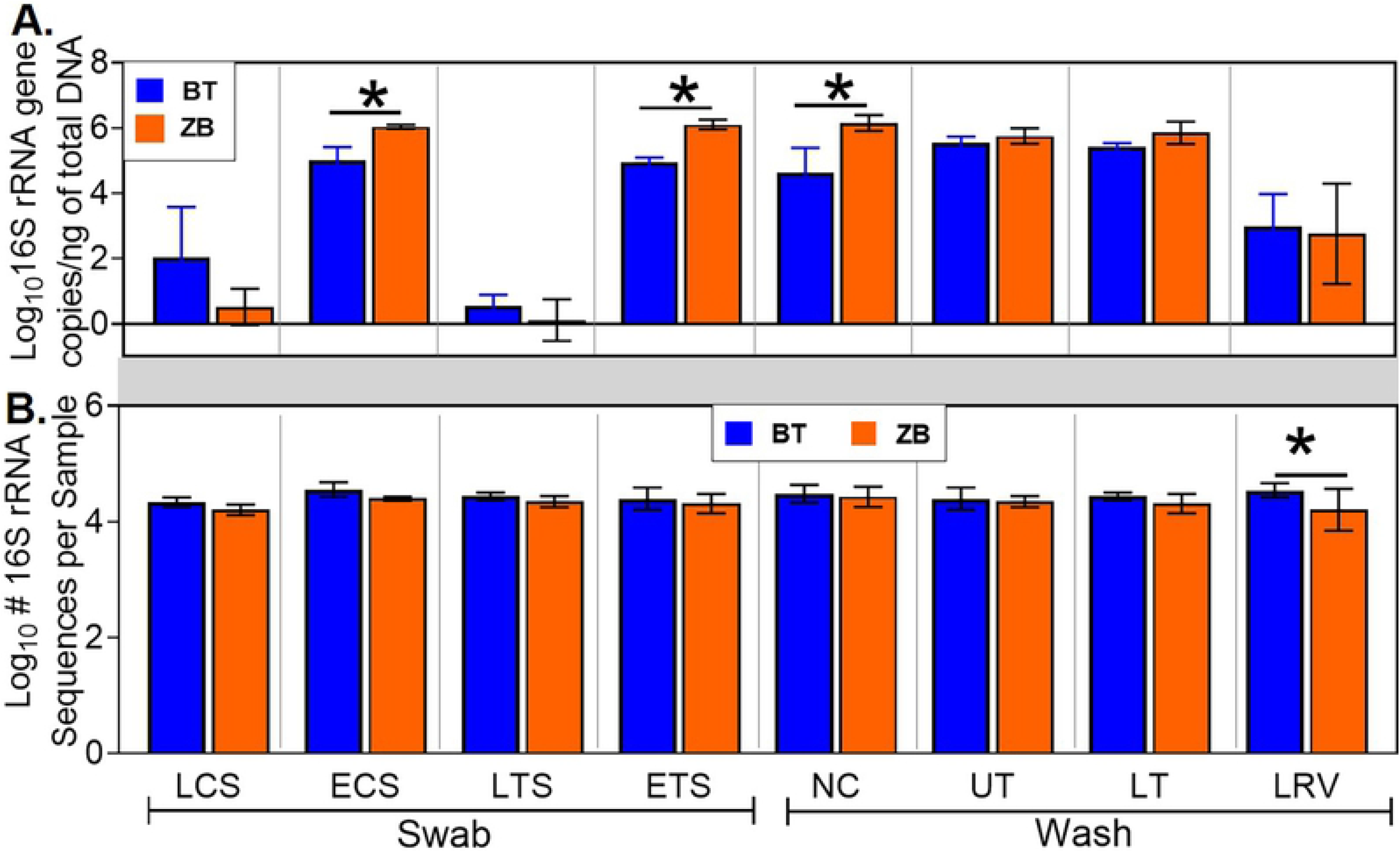
Between-kit comparison of pre- and post-sequencing 16S rRNA gene copies for each sample type. (A) Pre-sequencing 16S rRNA gene copies per nanogram of total DNA. Quantitative PCR was used to determine the number of 16S rRNA gene copies per μL of sample. The DNA samples were then normalized to 1.67 × 10^5^ 16S rRNA gene copies/μL for subsequent PCR prior to sequencing. (B) Post-sequencing 16S rRNA gene sequence counts. The sequences were quality-filtered as described in Materials and Methods. Statistical differences, * p < 0.05 (pairwise Kruskal-Wallis tests, with a Benjamini-Hochberg correction for false discovery). LCS, live choanal swab; ECS, euthanized choanal swab; LTS, live tracheal swab; ETS, euthanized tracheal swab; NC, nasal cavity wash; UT, upper trachea wash; LT, lower trachea wash; LRV, lower respiratory lavage.

### Minimal contaminants were identified, and the kits displayed subtle differences in extraction efficiencies

Several bacterial taxa were found in the negative extraction controls, indicating the presence of contaminants originating from either the reagents, sample processing steps, or sequence library preparation. These taxa were applied to a statistical classification procedure in R package decontam (version 1.2.1) [34] to identify contaminants in the respiratory samples. Of the 1487 ASVs detected among the respiratory samples, 14 were identified to be contaminants (Table 1). The contaminants were detected in low abundances (< 0.15 % relative abundance) independently of the kit used for extraction.

**Table 1.** Contaminants identified in Negative and Mock community controls

|  | Negative |  | Mock Community |  |
| --- | --- | --- | --- | --- |
|  | BT | ZB | BT | ZB |
| *Acetobacteroides* | 0.004* | 0.000 |  |  |
| *Acinetobacter* | 0.004 | 0.000 |  |  |
| *Bacillus* | 0.000 | 0.034 |  |  |
| *Bacteroides* |  |  | 0.011 | 0.000 |
| *Candidatus Woesebacteria* | 0.004 | 0.000 |  |  |
| Clostridiales | 0.004 | 0.006 |  |  |
| *Clostridium sensu stricto* 1 | 0.004 | 0.000 |  |  |
| *Corynebacterium* 1 | 0.020 | 0.138 |  |  |
| *Faecalibacterium* |  |  | 0.003 | 0.007 |
| *Flavobacterium* |  |  | 0.003 | 0.000 |
| *Halomonas* |  |  | 0.033 | 0.026 |
| *Lactobacillus* |  |  | 0.090 | 0.057 |
| *Lactobacillus vaginalis* |  |  | 0.000 | 0.026 |
| *Phascolarctobacterium* | 0.000 | 0.052 |  |  |
| *Phyllobacterium* | 0.004 | 0.006 |  |  |
| *Polymorphobacter* | 0.004 | 0.000 |  |  |
| *Pseudoalteromonas* |  |  | 0.003 | 0.000 |
| *Pseudochrobactrum* |  |  | 0.000 | 0.009 |
| *Ruminiclostridium* | 0.004 | 0.000 |  |  |
| Ruminococcaceae | 0.004 | 0.006 |  |  |
| *Sphingomonas* |  |  | 0.029 | 0.037 |
| *Stenotrophomonas acidaminiphila* | 0.004 | 0.000 |  |  |
| *Streptococcus* |  |  | 0.004 | 0.000 |
| *Streptococcus varani* |  |  | 0.000 | 0.004 |
| uncultured bacterium | 0.004 | 0.000 |  |  |
| *Vibrio* | 0.141 | 0.143 |  |  |
| **Total abundance** | **0.205** | **0.384** | **0.177** | **0.167** |

\*Values represent relative abundance as a percentage of pooled sequences from all sample types extracted with the indicated kit.

A mock community control was processed alongside the respiratory samples to assess the efficiency of extraction between the kits and the extent of contamination during the extraction process. Contaminants in the mock community samples were identified based on their absence in the compositional shotgun sequence data provided by ZymoBIOMICS in the certificate of analysis. Consistent with our previous findings [17], the mock community had < 1% contaminant bacteria (Table 1). While all bacteria in the mock community were detected in samples extracted with either kit, their relative abundances based on 16S rRNA sequencing varied compared to the expected abundances detected by shotgun sequencing (Table 2) [35]. Since the SILVA 132 classifier was not able to classify most of the bacteria in the mock community to the species level, our data does not account for the variability of 16S gene operons in each species. Therefore, it is impossible to directly compare the relative abundances observed in this study to the expected abundances based on shotgun sequencing. Nevertheless, the disparity in the relative abundance of 16S rRNA gene sequences between the kits may be indicative of differences in extraction efficiencies. The relative abundances of *Lactobacillus fermentum*, *Enterococcus*, *Enterobacteriaceae, Escherichia-Shigella,* and *Pseudomonas* were higher in samples extracted with the BT kit (Table 2). In contrast, the ZB kit was more efficient for the extraction of *Staphylococcus, Listeria,* and *Bacillus*.

**Table 2.** Efficiency of Mock community extraction

| Mock Community<br>Taxonomic Identity | Relative Abundance % |  |  |
| --- | --- | --- | --- |
|  | BT | ZB | Expected* |
| *Lactobacillus fermentum* | 25.6 | 21.8 | 11.7 |
| *Bacillus* | 15.6 | 16.3 | 12.3 |
| Enterobacteriaceae | 13.3 | 9.6 | 13.2 |
| *Escherichia-Shigella* | 13.2 | 10.2 | 13.6 |
| *Pseudomonas* | 12.6 | 9.0 | 12.9 |
| *Enterococcus* | 8.6 | 7.5 | 12.1 |
| *Staphylococcus* | 6.5 | 15.4 | 11.9 |
| *Listeria* | 4.3 | 9.9 | 12.2 |

\*Expected values were calculated based on shotgun sequencing as indicated in the certificate of analysis provided by ZymoBIOMICS for Catalog No: D6300; Lot No: ZRC190633
(https://s3.amazonaws.com/cofaman/cofaman/D6300_ZRC190633CofA.pdf).

### Method of extraction had minimal effects on taxonomic composition and diversity

After filtering the sequence data based on the 16S rRNA copy numbers and contaminants in negative extraction controls, the impacts of the extraction kits on basic microbiota parameters were assessed. Both of the extraction kits were able to detect the following predominant taxa at the class level: Bacilli, Clostridia, Gammaproteobacteria, Bacteroida, Erysipelotrichia, Actinobacteria, Mollicutes, and Alphaproteobacteria (Fig 4). Furthermore, several ASVs that were shared between the extraction methods within each sample type constituted > 92% of total abundance in the corresponding bacterial community (Table 3).

**Fig 4.**
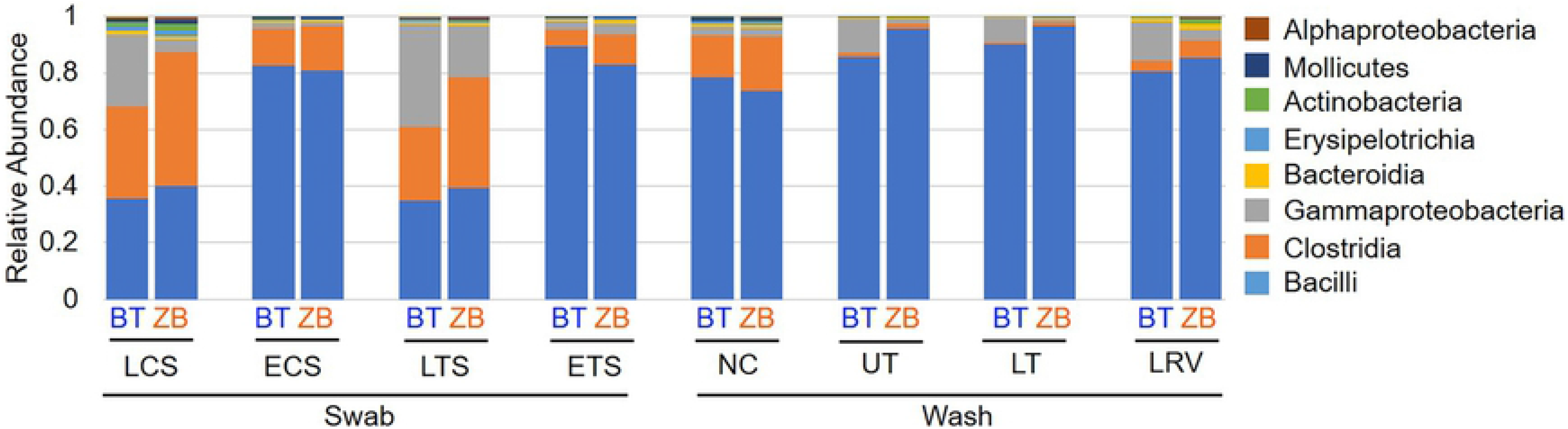
Comparison of relative abundances of different bacteria classes in samples extracted using the BT and ZB kits. LCS, live choanal swab; ECS, euthanized choanal swab; LTS, live tracheal swab; ETS, euthanized tracheal swab; NC, nasal cavity wash; UT, upper trachea wash; LT, lower trachea wash; LRV, lower respiratory lavage.

**Table 3.** Comparison of the number of shared ASVs and their relative abundance between the kits

| Sample type | # of shared ASVs | Relative Abundance (%) |  |
| --- | --- | --- | --- |
|  |  | BT | ZB |
| Live tracheal Swab | 103 | 92.1 | 97.8 |
| Euthanized tracheal swab | 99 | 99.5 | 99.5 |
| Upper tracheal wash | 59 | 97.7 | 99.5 |
| Lower tracheal wash | 46 | 98.6 | 99.6 |
| Live choanal swab | 167 | 95.2 | 98.6 |
| Euthanized choanal swab | 104 | 99.4 | 99.7 |
| Nasal cavity Wash | 155 | 99.6 | 99.7 |
| Lower respiratory lavage | 135 | 98.4 | 96.9 |

The characteristics of the microbial communities detected by each kit were evaluated further through alpha and beta diversity metrics. In majority of the samples, the number of observed ASVs in a given sample type were statistically indistinguishable between the kits, except in the live choanal swab where, on average, samples extracted using the BT kit had a higher number of ASVs (Fig 5A). Similarly, the extraction kit did not affect the species evenness in any of the sample types (Fig 5B). Beta diversity was assessed by comparing within sample type UniFrac distances between kits. No statistical differences were observed between the kits (Fig 6).

**Fig 5.**
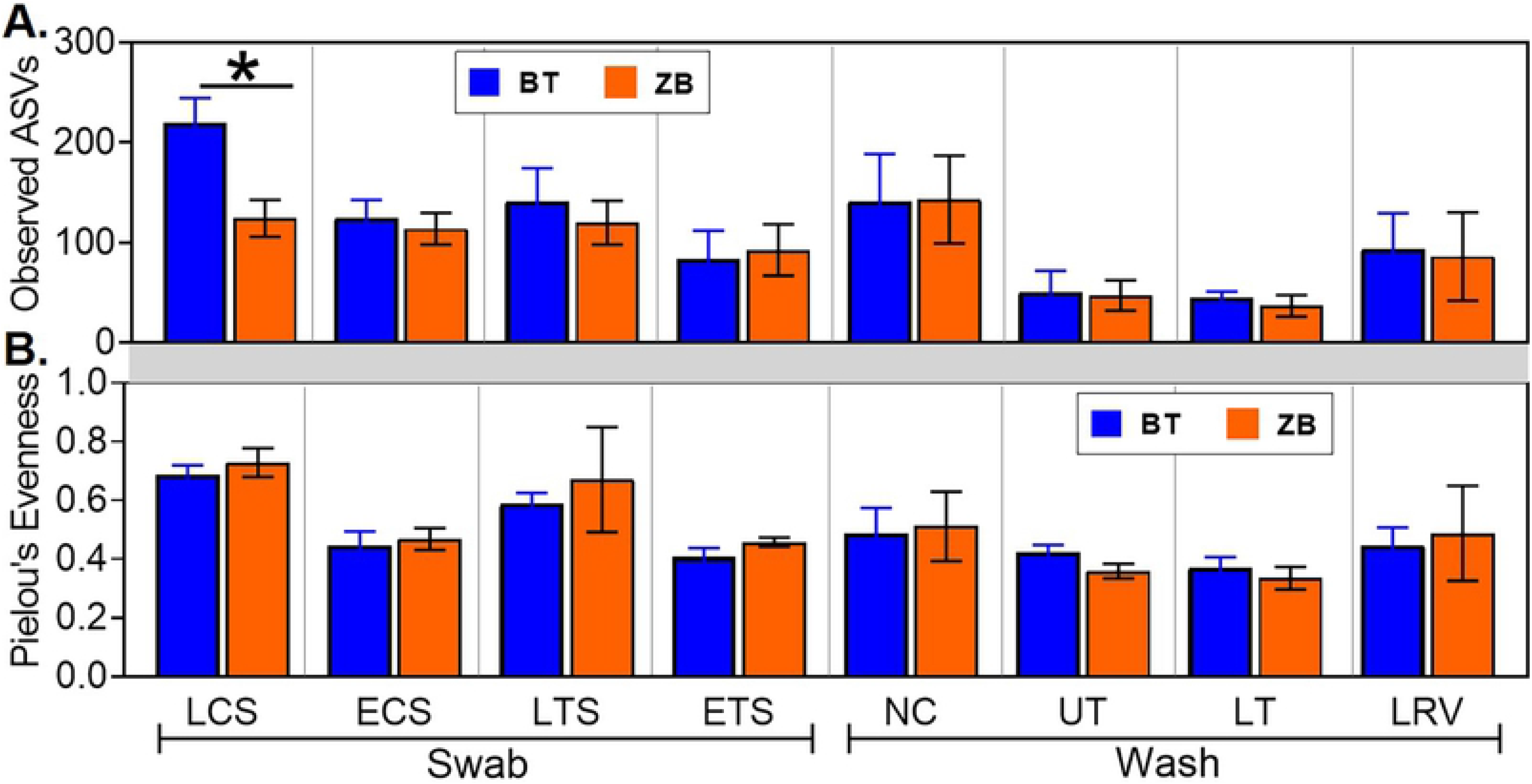
Comparison of alpha diversity metrics from samples extracted using the ZB and BT kits. (A) Species richness was estimated based on the number of uniquely observed amplicon sequence variants (ASVs) in each sample. (B) Species evenness in each sample was estimated using Pielou’s evenness index. Statistical differences, * p < 0.05 (pairwise Kruskal-Wallis tests, with a Benjamini-Hochberg correction for false discovery). LCS, live choanal swab; ECS, euthanized choanal swab; LTS, live tracheal swab; ETS, euthanized tracheal swab; NC, nasal cavity wash; UT, upper trachea wash; LT, lower trachea wash; LRV, lower respiratory lavage.

**Fig 6.**
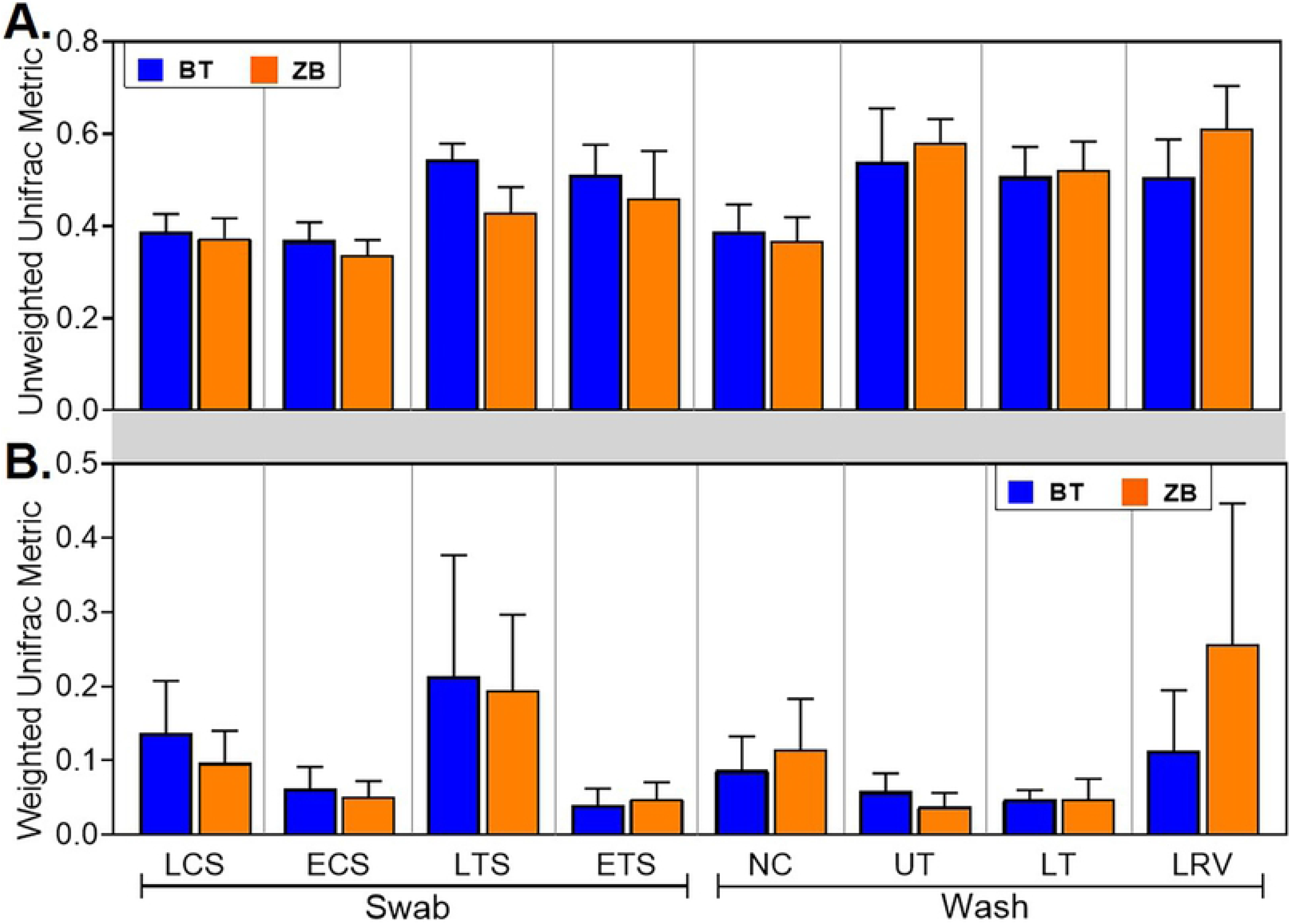
Comparison of beta diversity metrics from samples extracted using the BT and ZB kits. Unweighted (A) and Weighted (B) UniFrac distances were compared between samples from a given sample type. No statistical differences were found between the kits (pairwise permutational multivariate analysis of variance-PERMANOVA with a Benjamini-Hochberg correction for false discovery). LCS, live choanal swab; ECS, euthanized choanal swab; LTS, live tracheal swab; ETS, euthanized tracheal swab; NC, nasal cavity wash; UT, upper trachea wash; LT, lower trachea wash; LRV, lower respiratory lavage.

## Discussion

The DNA extraction process can be a major source of variation between microbiome studies [7]. This poses a potential problem with inter-study comparison among the growing number of poultry respiratory microbiome studies. Using the BT kit in our previous studies, we were able to characterize the microbial communities present in sinus cavities, tracheas and lower respiratory tracts of turkeys, chicken broilers, and chicken layers raised in commercial settings, and specific-pathogen free chickens raised for research [2, 15–17]. Although we optimized the BT kit for increased DNA yields and removal of PCR inhibitors from respiratory samples [2, 15, 17], the kit is not specifically certified for microbiome sample extraction [29]. Here, we compared the performance of the BT kit with that of the ZB kit, which is validated for extraction of low-bioburden microbiome samples [30]. Overall, while statistical differences were observed between kits in the pre-sequencing results, post-sequencing microbiota data obtained with normalized samples showed only minimal differences between the kits.

Both kits were very efficient in eliminating protein contaminants absorbing at 280 nm as indicated by A260/A280 ratios within the expected range of 1.8 – 2.0 [29, 37, 38]. However, the ZB kit seemed inefficient at removing non-protein contaminants absorbing at 230 nm from some sample types, especially the nasal cavity wash and lower respiratory lavage samples, as indicated by A260/A230 ratios that were below the expected range of 2.2-2.5 (Fig 2B) [38]. While the exact nature of these impurities remain undetermined, their impact on PCR amplification was negligible as samples extracted with the ZB kit tended to have similar or higher 16S rRNA gene copy numbers compared to those extracted with the BT kit (Fig 3A). Nevertheless, the ZB kit may require further optimization since the A260/A230 ratios reported in other studies comparing the performances of several commercial kits for bacterial DNA extraction from different sample types were all within the expected range [22, 40–41].

Our protocol for 16S rRNA sequencing on the Illumina platform calls for sample normalization to 5 × 10^5^ 16S rRNA copies/ 3μL before PCR amplification and library preparation to have a 10X target sequencing coverage. This normalization appeared to eliminate between-kit differences in the number of 16S rRNA sequences (Fig 3B). Studies evaluating kit effects on bacterial diversity typically focus on the differences observed in the concentration of 16S rRNA copies detected via quantitative PCR without paying much attention to the actual number of 16S rRNA sequences recovered [40, 42]. Furthermore, the contribution of unnormalized DNA concentrations to the observed microbiota profile differences between kits needs to be clarified.

In addition to pre-sequencing data, bacterial compositions and diversity metrics are commonly used as proxies for performance in extraction kit-comparison studies [22, 41, 43–44]. Here, all the predominant ASVs detected in samples extracted with one kit were present in replicate samples extracted with the other kit and did not show great differences at the class level (Table 3 and Fig 4). Similarly, between-kits differences in alpha and beta diversity metrics were statistically indistinguishable (Figs 5-6).

Other studies have typically compared multiple extraction kits using a single sample type [22, 41, 43–44]. In this study, although post-sequencing analysis revealed comparable performances between the BT and ZB kits across multiple types of respiratory samples, the impact of sample type on other DNA extraction methods requires further investigation. For example, between-kit differences appear to be magnified in gut samples [40, 45] and reduced in respiratory samples [42, 43]. Nevertheless, both BT and ZB kits performed similarly in terms of extracting specific bacteria in the mock community and showed differences in bacterial compositions and diversity metrics between different respiratory sites (Tables 2 and 3), and have therefore validated our previous reports [2, 15–17]. With that said, the ZB kit is less likely to contain contaminant bacterial DNA, thereby reducing the likelihood of excluding samples with low DNA concentrations from analysis.

The design of this study resulted in uneven distribution of sample replicates between sample types. For each kit, there were 8 replicates/sample type for lavage and nasal wash samples and 4 replicates/sample type for tracheal wash and swab samples. Therefore, reasonable comparisons between sample types within each kit could not be achieved in this study.

In conclusion, both the ZB and BT methods of DNA extraction produced similar chicken respiratory microbiome data in terms of bacterial diversity and class-level taxonomic profiles.

## Acknowledgments

The authors would like to acknowledge the professional and enthusiastic support from Dr. Juliette Hanson and Megan Strother for animal welfare. Sequence processing and analysis were performed using the resources of the Molecular & Cellular Imaging Center, Ohio Agricultural Research and Development Center, The Ohio State University. This project was supported by Agriculture and Food Research Initiative competitive grant number 2015-68004-23131 from the USDA National Institute of Food and Agriculture (to C.-W.L.).

## REFERENCES

1. Simon K, Verwoolde MB, Zhang J, Smidt H, de Vries Reilingh G, Kemp B, et al. Long-term effects of early life microbiota disturbance on adaptive immunity in laying hens. Poult Sci. 2016;95: 1543–1554. doi:10.3382/ps/pew088

2. Johnson TJ, Youmans BP, Noll S, Cardona C, Evans NP, Karnezos TP, et al. A Consistent and Predictable Commercial Broiler Chicken Bacterial Microbiota in Antibiotic-Free Production Displays Strong Correlations with Performance. Appl Environ Microbiol. 2018;84. doi:10.1128/AEM.00362-18

3. Torok VA, Hughes RJ, Mikkelsen LL, Perez-Maldonado R, Balding K, MacAlpine R, et al. Identification and Characterization of Potential Performance-Related Gut Microbiotas in Broiler Chickens across Various Feeding Trials. Appl Environ Microbiol. 2011;77: 5868–5878. doi:10.1128/AEM.00165-11

4. Nava GM, Bielke LR, Callaway TR, Castañeda MP. Probiotic alternatives to reduce gastrointestinal infections: the poultry experience. Anim Health Res Rev. 2005;6: 105–118. doi:10.1079/ahr2005103

5. Stanley D, Denman SE, Hughes RJ, Geier MS, Crowley TM, Chen H, et al. Intestinal microbiota associated with differential feed conversion efficiency in chickens. Appl Microbiol Biotechnol. 2012;96: 1361–1369. doi:10.1007/s00253-011-3847-5

6. Costea PI, Zeller G, Sunagawa S, Pelletier E, Alberti A, Levenez F, et al. Towards standards for human fecal sample processing in metagenomic studies. Nat Biotechnol. 2017;35: 1069–1076. doi:10.1038/nbt.3960

7. Sinha R, Abu-Ali G, Vogtmann E, Fodor AA, Ren B, Amir A, et al. Assessment of variation in microbial community amplicon sequencing by the Microbiome Quality Control (MBQC) project consortium. Nature Biotechnology. 2017;35: 1077–1086. doi:10.1038/nbt.3981

8. Koren O, Knights D, Gonzalez A, Waldron L, Segata N, Knight R, et al. A Guide to Enterotypes across the Human Body: Meta-Analysis of Microbial Community Structures in Human Microbiome Datasets. PLOS Computational Biology. 2013;9: e1002863. doi:10.1371/journal.pcbi.1002863

9. Brooks JP, Edwards DJ, Harwich MD, Rivera MC, Fettweis JM, Serrano MG, et al. The truth about metagenomics: quantifying and counteracting bias in 16S rRNA studies. BMC Microbiol. 2015;15: 66. doi:10.1186/s12866-015-0351-6

10. Schirmer M, Ijaz UZ, D’Amore R, Hall N, Sloan WT, Quince C. Insight into biases and sequencing errors for amplicon sequencing with the Illumina MiSeq platform. Nucleic Acids Res. 2015;43: e37. doi:10.1093/nar/gku1341

11. Bag S, Saha B, Mehta O, Anbumani D, Kumar N, Dayal M, et al. An Improved Method for High Quality Metagenomics DNA Extraction from Human and Environmental Samples. Sci Rep. 2016;6: 26775. doi:10.1038/srep26775

12. Baker GC, Smith JJ, Cowan DA. Review and re-analysis of domain-specific 16S primers. J Microbiol Methods. 2003;55: 541–555. doi:10.1016/j.mimet.2003.08.009

13. Cruaud P, Vigneron A, Lucchetti-Miganeh C, Ciron PE, Godfroy A, Cambon-Bonavita M-A. Influence of DNA Extraction Method, 16S rRNA Targeted Hypervariable Regions, and Sample Origin on Microbial Diversity Detected by 454 Pyrosequencing in Marine Chemosynthetic Ecosystems. Appl Environ Microbiol. 2014;80: 4626–4639. doi:10.1128/AEM.00592-14

14. Tremblay J, Singh K, Fern A, Kirton ES, He S, Woyke T, et al. Primer and platform effects on 16S rRNA tag sequencing. Front Microbiol. 2015;6. doi:10.3389/fmicb.2015.00771

15. Ngunjiri JM, Taylor KJM, Abundo MC, Jang H, Elaish M, Kc M, et al. Farm Stage, Bird Age, and Body Site Dominantly Affect the Quantity, Taxonomic Composition, and Dynamics of Respiratory and Gut Microbiota of Commercial Layer Chickens. Appl Environ Microbiol. 2019;85. doi:10.1128/AEM.03137-18

16. Taylor KJM, Ngunjiri JM, Abundo MC, Jang H, Elaish M, Ghorbani A, et al. Respiratory and Gut Microbiota in Commercial Turkey Flocks with Disparate Weight Gain Trajectories Display Differential Compositional Dynamics. Appl Environ Microbiol. 2020;86. doi:10.1128/AEM.00431-20

17. Abundo MC, Ngunjiri JM, Taylor KJM, Ji H, Ghorbani A, K C M, et al. Evaluation of Sampling Methods for the Study of Avian Respiratory Microbiota. Avian Diseases. 2020;In Press. doi:10.1637/aviandiseases-D-19-00200

18. Vermassen A, Leroy S, Talon R, Provot C, Popowska M, Desvaux M. Cell Wall Hydrolases in Bacteria: Insight on the Diversity of Cell Wall Amidases, Glycosidases and Peptidases Toward Peptidoglycan. Front Microbiol. 2019;10. doi:10.3389/fmicb.2019.00331

19. Shehadul Islam M, Aryasomayajula A, Selvaganapathy PR. A Review on Macroscale and Microscale Cell Lysis Methods. Micromachines (Basel). 2017;8. doi:10.3390/mi8030083

20. Yu Z, Morrison M. Improved extraction of PCR-quality community DNA from digesta and fecal samples. BioTechniques. 2004;36: 808–812. doi:10.2144/04365ST04

21. Liu Z, Lozupone C, Hamady M, Bushman FD, Knight R. Short pyrosequencing reads suffice for accurate microbial community analysis. Nucleic Acids Res. 2007;35: e120. doi:10.1093/nar/gkm541

22. Sohrabi M, Nair RG, Samaranayake LP, Zhang L, Zulfiker AHM, Ahmetagic A, et al. The yield and quality of cellular and bacterial DNA extracts from human oral rinse samples are variably affected by the cell lysis methodology. J Microbiol Methods. 2016;122: 64–72. doi:10.1016/j.mimet.2016.01.013

23. Salter SJ, Cox MJ, Turek EM, Calus ST, Cookson WO, Moffatt MF, et al. Reagent and laboratory contamination can critically impact sequence-based microbiome analyses. BMC Biology. 2014;12: 87. doi:10.1186/s12915-014-0087-z

24. Karstens L, Asquith M, Davin S, Fair D, Gregory WT, Wolfe AJ, et al. Controlling for Contaminants in Low-Biomass 16S rRNA Gene Sequencing Experiments. mSystems. 2019;4. doi:10.1128/mSystems.00290-19

25. Shabbir MZ, Malys T, Ivanov YV, Park J, Shabbir MAB, Rabbani M, et al. Microbial communities present in the lower respiratory tract of clinically healthy birds in Pakistan. Poult Sci. 2015;94: 612–620. doi:10.3382/ps/pev010

26. Glendinning L, McLachlan G, Vervelde L. Age-related differences in the respiratory microbiota of chickens. PLoS One. 2017;12. doi:10.1371/journal.pone.0188455

27. Sohail MU, Hume ME, Byrd JA, Nisbet DJ, Shabbir MZ, Ijaz A, et al. Molecular analysis of the caecal and tracheal microbiome of heat-stressed broilers supplemented with prebiotic and probiotic. Avian Pathol. 2015;44: 67–74. doi:10.1080/03079457.2015.1004622

28. Kursa O, Tomczyk G, Sawicka-Durkalec A, Giza A, Słomiany-Szwarc M. Characterization of the upper respiratory tract microbiome of turkeys. In Review; 2020 Jun. doi:10.21203/rs.3.rs-33858/v1

29. DNeasy® Blood & Tissue Handbook July 2020. https://www.qiagen.com/us/resources/download.aspx?id=68f29296-5a9f-40fa-8b3d-1c148d0b3030&lang=en 2020 [cited 27 Aug 2020]. Available: https://www.qiagen.com/us/resources/download.aspx?id=68f29296-5a9f-40fa-8b3d-1c148d0b3030&lang=en

30. ZymoBIOMICS™ DNA Miniprep Kit Instruction Manual ver 1.4.1. 27 Aug 2020 [cited 27 Aug 2020]. Available: https://files.zymoresearch.com/protocols/_d4300t_d4300_d4304_zymobiomics_dna_miniprep_kit.pdf

31. Bolyen E, Rideout JR, Dillon MR, Bokulich NA, Abnet CC, Al-Ghalith GA, et al. Reproducible, interactive, scalable and extensible microbiome data science using QIIME 2. Nature Biotechnology. 2019;37: 852–857. doi:10.1038/s41587-019-0209-9

32. Nahm FS. Nonparametric statistical tests for the continuous data: the basic concept and the practical use. Korean J Anesthesiol. 2016;69: 8–14. doi:10.4097/kjae.2016.69.1.8

33. Anderson MJ. Permutational Multivariate Analysis of Variance (PERMANOVA). Wiley StatsRef: Statistics Reference Online. American Cancer Society; 2017. pp. 1–15. doi:10.1002/9781118445112.stat07841

34. Davis NM, Proctor DM, Holmes SP, Relman DA, Callahan BJ. Simple statistical identification and removal of contaminant sequences in marker-gene and metagenomics data. Microbiome. 2018;6: 226. doi:10.1186/s40168-018-0605-2

35. Zymo Research. Certificate of Analysis. 21 Aug 2020 [cited 21 Aug 2020]. Available: https://s3.amazonaws.com/cofaman/cofaman/D6300_ZRC190633CofA.pdf

36. Koetsier G, Cantor E. A Practical Guide to Analyzing Nucleic Acid Concentration and Purity with Microvolume Spectrophotometers. New England BioLabs: Technical note. 2019; 8.

37. Glasel J. Validity of Nucleic Acid Purities Monitored by A260/A280 Absorbance Ratios. BioTechniques. 1995; 62–63.

38. Laws GM, Adams SP. Measurement of 8-OHdG in DNA by HPLC/ECD: The Importance of DNA Purity. BioTechniques. 1996;20: 36–38. doi:10.2144/96201bm06

39. Pankoke H, Maus I, Loh G, Hüser A, Seifert J, Tilker A, et al. F5Evaluation of commercially available DNA extraction kits for the analysis of the broiler chicken cecal microbiota. FEMS Microbiology Letters. 2019 [cited 28 Aug 2020]. doi:10.1093/femsle/fnz033

40. Teng F, Darveekaran Nair SS, Zhu P, Li S, Huang S, Li X, et al. Impact of DNA extraction method and targeted 16S-rRNA hypervariable region on oral microbiota profiling. Scientific Reports. 2018;8: 16321. doi:10.1038/s41598-018-34294-x

41. Pérez-Losada M, Crandall KA, Freishtat RJ. Comparison of two commercial DNA extraction kits for the analysis of nasopharyngeal bacterial communities. microbiology 2016, Vol 2, Pages 108–119. 2016 [cited 29 Jun 2020]. doi:10.3934/microbiol.2016.2.108

42. Terranova L, Oriano M, Teri A, Ruggiero L, Tafuro C, Marchisio P, et al. How to Process Sputum Samples and Extract Bacterial DNA for Microbiota Analysis. Int J Mol Sci. 2018;19. doi:10.3390/ijms19103256

43. Lazarevic V, Gaïa N, Girard M, François P, Schrenzel J. Comparison of DNA Extraction Methods in Analysis of Salivary Bacterial Communities. PLOS ONE. 2013;8: e67699. doi:10.1371/journal.pone.0067699

44. Lim MY, Song E-J, Kim SH, Lee J, Nam Y-D. Comparison of DNA extraction methods for human gut microbial community profiling. Systematic and Applied Microbiology. 2018;41: 151–157. doi:10.1016/j.syapm.2017.11.008

